# Spatial analysis of American Eel (*Anguilla rostrata*), fish passage and land use in Chesapeake Bay tributaries

**DOI:** 10.1101/2020.03.17.995183

**Authors:** Nicholas J. Walker, V. Prasad, K. de Mutsert, C.A. Dolloff, A.A. Aguirre

**Author notes:** These authors contributed equally to this work.

## Abstract

Catadromous eels are found in more habitats than any other fish and are capable of inhabiting marine, brackish and freshwater environments. In this study we used the American Eel (*Anguilla rostrata*) as a bioindicator organism to create a novel method of using spatial analysis to study species conservation over landscape scales. We built a model of the subwatersheds of the Chesapeake Bay using a Digital Elevation Model (DEM) and overlaid eel density data (> 1 million eels sampled), dam density data and land use in ArcGIS. Dam construction in the study area peaked between 1955 and 1975, possibly as a result of flood control measures. Effects of land use were localized and most pronounced in areas around Baltimore, Maryland, Washington, D.C. and Richmond, Virginia, USA. Results indicate the Potomac and Rappahannock rivers appear to be areas of lesser concern while the upper James and York rivers are ideal for follow-up studies, since these area rank poorly in both eel density and barriers to fish passage. Because these rivers have high eel density downstream, the dams appear to be the limiting factor. Sampling methods have been inconsistent over time, making it is difficult to determine where eel densities are low vs. the area having had little sampling effort. This is partially resolved with catch per sampling event (CPSE), which appears to show a relationship between eels sampled and the number caught per sample. Potential strategies for improving watersheds include dam removal, fish passage and habitat restoration.

## Introduction

The American Eel (*Anguilla rostrata*) is a catadromous species that spawns in the Sargasso Sea and is distributed by ocean currents to North America, the Caribbean and the northern drainages of South America [1]. This species can use one of several life strategies and live in saltwater, brackish estuarine water and freshwater [2, 3]. Catadromous eels are found in more diverse habitats than any other fish because of their adaptability [4], which makes them ideal candidates for studies over a large geographic region. Few studies however have studied eels at landscape scales taking into account entire watersheds and barriers to fish passage. Eels have been used as sentinel species for studying environmental contamination in streams [5] as well as in classroom studies combining biology and geography [6] and citizen science projects [7].

The species has been assessed as depleted in recent years [10, 11] although the decline was primarily during the latter half of the 20^th^ century and has leveled off in the last couple of decades [12]. Stressors include climate change, changes in the North Atlantic currents, introduced parasites, especially *Anguillocoloides crassus*, habitat loss, pollution, overfishing and physical barriers [9].

One of the challenges for American Eel is to be able to migrate upstream into non-tidal habitats (during the yellow phase) and then back downstream during the silver phase. Estuarine habitats used by American Eels have been lost because of filling and conversion to upland, as well as eutrophication and contaminants. Eels are able to climb very well for fish and can even travel up Great Falls on the Potomac River -as evidenced by their presence in upstream tributaries, including the Shenandoah River [9].

While there are still nine active silver eel weirs in the Delaware River and tributaries [20], all of the others have closed, with the Maine silver eel fishery closing in 2014 [21]. Thus, it appears the decline in American Eel is coincident with the construction of dams (which accelerated during the 1950s to 1970s) and the continued presence of these barriers could be the primary reason the population is still at around half of historical levels. Because of the large distribution of American Eel, future studies will need to take into account the entire range to determine the factors affecting carrying capacity.

American Eels can adapt to a variety of habitats through local adaptation and phenotype plasticity. The three life strategies (freshwater, brackish water and marine water) are tied to distinct ecotypes that can be consistently differentiated by polygenic genetic differences and blindly assigned to their habitat of origin, although the mechanism for how this occurs is unclear. Thus, it is necessary to conserve habitat and connectivity across the range to conserve genetic diversity [22]. Some of the historical range has already been reduced by the construction of dams for hydropower and water storage. Habitat loss from barriers is considered a historical effect and its population level effects have likely already been realized [23].

Throughout the freshwater habitats used by American Eel, rivers and streams are lined with varying degrees of riparian buffers and urbanized, impervious surfaces. Riparian buffers include both forest and grassland buffers, along with shrubs and other vegetation. These are managed to maintain the integrity of stream banks, reduce the impact of terrestrial pollution and to supply food and thermal protection to fish and other wildlife [36]. Riparian buffers can shade riverbeds, sustain allochthonous inputs (both of which reduce algal growth), intercept and absorb nutrients and support diverse habitats for fish communities. Buffers can also provide inputs of leaf litter and terrestrial invertebrates, which provide food for aquatic fauna [37]. Urbanization can affect shading, allochthonous inputs, hydrology, water chemistry (by altering stream geomorphology) water quality and invertebrate prey densities [14]. The relationship between eels and land use is complicated and cannot be taken as simply a causation between more land use buffers and increased eel densities.

We are aware of no previous studies examining the effects of land use buffers on American Eel in the Chesapeake Bay watershed, although this has been studied in the Hudson River estuary in New York. In this watershed, American Eel abundance and biomass respond strongly to barriers and secondarily to local-scale urbanization in tributary subcatchments [14]. At low levels, urbanization can result in higher eel abundance, but as it increases there are negative effects because of pollution and eutrophication, with sites at 30-40% urbanization having no eels at all [14]. Land use can also affect parasite incidence. Infection rates of *Eustrongylides tubifex* and *A. crassus* in the Hudson River increased with urbanized land, as well as higher water temperatures [38]. In this estuary, eels of all sizes use leaf litter as cover during autumn and substrate cover during summer. As such, protecting riparian forest buffers can help provide deciduous leaf cover for eels in autumn [39].

The effect of riparian buffers on eels has also been studied in New Zealand, on both longfin eels (*A. dieffenbachii*) and shortfin eels (*A. australis*). The abundance of both species is associated with increased riparian cover [40]. There are differences in habitat selection between the two species, but this would not appear to be a factor in North America since there is only one eel species here. In New Zealand, eel total length is associated with riparian habitats. In a short-term case study in New Zealand, removal of overhanging vegetation and in-stream wood from short reaches of a small pastoral stream with intact riparian margins resulted in the formation of shallow uniform runs rather than the pool and riffle structures found in unmodified reaches. This reduced the abundance of adult longfin eel (*A. dieffenbachii*) although elvers became more abundant [41]. If these results apply to the Chesapeake Bay, the loss of upstream riparian buffers could result in reduced yellow eel abundance, since elvers are only found at the mouth of the bay.

In some studies however, eel presence has been negatively correlated with native New Zealand forests, with greater eel biomass and a density along pastoral sites [42]. Perhaps paradoxically, eel abundance and biomass are correlated with non-native willow trees (Glova 1994). Eels are largest in areas with pasture, medium in areas with willows and smallest along tussock (grasslands) [41].

The objective of this research was to create a model of eel migration and habitat suitability over a large geographic area by combining biological data on eels with environmental data and a Digital Elevation Model (DEM) of the watershed. DEMs have been used in spatial analysis to construct drainage paths for decades [43, 44] and our goal was to create an “eel’s eye view” of the Chesapeake Bay for migratory eels. This is the first study to combine over a century of eel abundance and density data across two dozen Chesapeake Bay tributaries along with dams and land use to determine habitat suitability and priority areas for conservation measures. The goal was to create a new model for species conservation using spatial analysis.

## Materials and methods

A map of the Chesapeake Bay was created using ESRI ArcGIS 10.6., to analyze several factors: eel density (and catch per sampling event -CPSE), presence of dams (with and without fish passage) and land use around streams. The eel data comes from a database we compiled using datasets from the Virginia Department of Game and Inland Fisheries (including the JFISH collection and data originally collected by the Smithsonian Institution), Maryland Department of Natural Resources (including the Maryland Biology Stream Survey, MDCHES database and SASSFish Index), the U.S. Forest Service Southern Research Station and the U.S. Fish & Wildlife Service. The dataset is described in a separate paper (Walker et al. in review).

In this work, we used location, count, sampling events and life stage. Although eel data was available from 1911 to 2018, it was sparse prior to 2000. This limits the ability to study the eel population prior to the onset of commercial fishing, 1952 [45] or the start of the decline two decades later [46]. Other limitations included the type of fishing gear used was only available in about 1/4 of the records and varied between backpack electrofishing, boat electrofishing, eel pots and Irish eel ramps; hence fishing gear was not included in this analysis. The study area was comprised of Virginia, Maryland and Washington, D.C.

The database of dams was provided by The Nature Conservancy [47]. There are 3,828 dams in the Chesapeake Bay watershed. The Nature Conservancy provided additional data on dams, including which ones have fish passage provisions (Erik Martin, pers. comm. YEAR). Land use data for the year 2015 was provided by the Université Catholique de Louvain [48].

Delineating locations as “land” and “water” required a multi-step process. First, using the ArcGIS tool *Buffer* (Analysis) a 100 km circle was drawn around all eel sample points, with dissolve type set to “All”. The “Water Courses – Global Map” shapefile from the USGS was clipped to this buffer using *Clip* (Analysis) and the resulting attribute table was exported to Microsoft Excel format. We created a spreadsheet and traced each segment back to the mainstem of the rivers flowing into the Bay. This made it apparent that some of the eels sampled (approximately 15%) had not come through the Chesapeake Bay, but through other drainages, primarily the Roanoke and Monongahela Rivers. These were excluded from analysis.

To determine the extent of the rivers, we combined four data sources. The first two were from the USGS National Map: Small Scale (https://nationalmap.gov/small_scale/atlasftp.html) [49]. We used “Streams, One Million-Scale” and “Water Courses – Global Map”, both at 1:1,000,000 scale (100 m). Additionally, we created a map of streams using 20 tiles from the ASTER Global Digital Elevation Model (DEM) dataset with a resolution of 1-arc-second (30×30xm) [50]. We used the following Spatial Analyst tools in ArcGIS to generate all streams > 25 km^2^: Fill, Flow Direction and Flow Accumulation, followed by the Raster Calculator with *SetNull(“bay_flowac” < 27778,1)* to set the minimum threshold to 25 km^2^ (25*10^6^)/(30^2^), then Stream Link, Stream Order and Stream to Feature. The two USGS stream files were merged with each other but could not be merged with the streams we generated from the DEM, because of differences in the data formats in the attribute tables. Therefore, a one km buffer (referring to the *Buffer* (Analysis) tool in ArcGIS, not a riparian or land use buffer) was drawn around the USGS streams and a separate one-km ArcGIS *Buffer* was drawn around the DEM-created streams (using the Dissolve (All) option in both cases) and the resulting polygons were merged. The areas marked as water on the land use raster were converted to a shapefile and a 1 km ArcGIS *Buffer* was applied. This was merged with the preceding polygons and another one-km ArcGIS *Buffer* was used to remove a few gaps and to make the streams more visible when the map is zoomed out. Thus, there is approximately a two-km polygon around each polyline marked as water. The ArcGIS *Buffers* are necessary because a polyline is a vector with no width and does not reflect the width of streams and creeks in real life.

The ASTER data was used to generate the watersheds via a separate process, using the following Spatial Analyst tools: Fill, Flow Direction, Basin and Raster to Vector. This generated a map with over 300,000 polygons. Polygons that coincided with the river segments were merged using the ArcGIS editor, all other polygons were removed and the remaining polygons were joined to remove the grid created by the initial 20 raster tiles. The resulting polygons each represent watersheds and are hereafter referred to as the study area. Watersheds were further divided into subwatersheds at approximately 50 km along the mainstem. Any watershed with less than 50 km on the mainstem was not subdivided. These subwatersheds were used to calculate the densities of eels and dams and bin the results into categories for ranking.

The generated watersheds do not include the full extent of several rivers for two reasons: 1) the tasks become exponentially more computationally difficult with increased area, especially at high resolution; and 2) we wanted to focus our efforts on the areas where we had eel data, which were Virginia, Maryland and Washington D.C. We removed sections of Delaware, Pennsylvania and West Virginia when mapping the watersheds and did not include the excess areas when calculating eel and dam densities. We believe this is not a significant limitation to this study because the areas closest to the mouths of the rivers are most important for studying the habitats and passage of catadromous eels. The state boundaries were clipped after the watersheds had already been generated based on DEM and thus some sections (most notably Potomac River 05) are split in half by the boundaries, but the two fragments are processed together in the model because they are still part of the same subwatershed section.

When the eel data were placed on the base map, it became apparent that much of it did not align with conventional maps of streams and creeks, with eels appearing to be on land rather than water. This may be due to errors in data collection, or eels that were caught from smaller creeks not visible on most maps, or ephemeral streams, or even streams that have changed course since the data collection. In some cases, the coordinates may be slightly off, the result of over a century of data collection using a variety of methods, formats and record keeping. Our methodology takes all of this into account by assigning each eel sample (point data) to a subwatershed that drains to specific pour points at the mouth of the Chesapeake Bay. This way, whether the fish appear to be on land or water, the model can determine the route these eels used to arrive at that point and what kind of barriers they may have encountered along their way.

In many cases the eels that appear to be over land actually are in small streams and creeks that do not appear on the map. For example, consider two eels from tributaries on the York River appear to have been caught over land (Fig. 1A). When these locations are analyzed on Google Earth, they align with small streams (< 10 m wide). These streams are not visible in the most recent Google Earth imagery from May 2018 (Fig. 1B) but can be seen on earlier imagery from April 2013 (Fig. 1C-D).

**Fig. 1.**
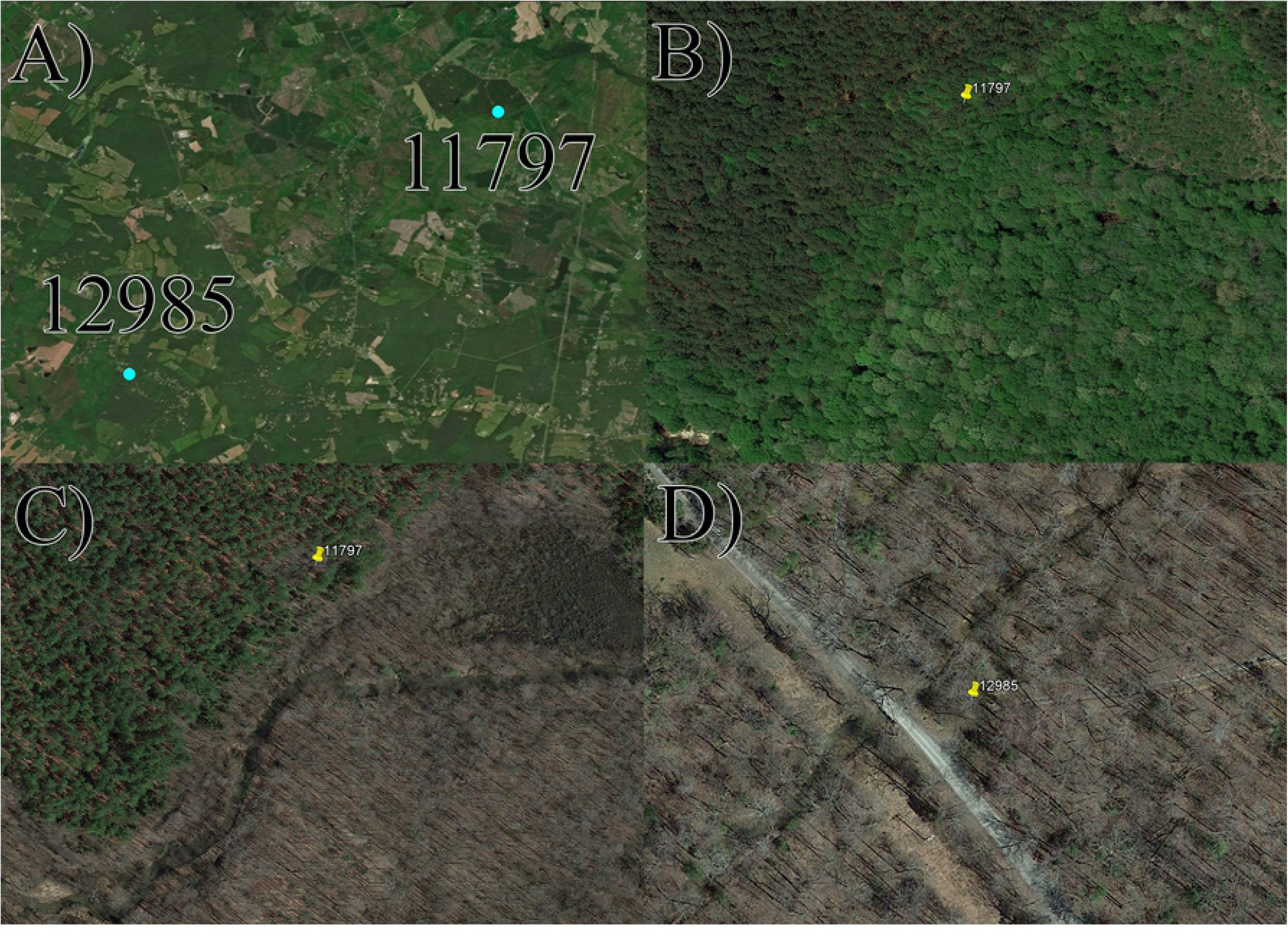
Eel locations coincident with small streams not visible on the stream map. 1A shows both locations (on tributaries of the York River) on the ArcGIS map. Location 11797 (37.926093 N, 77.801595 W) indicates three eels caught at Long Creek along Route 665 in Virginia on August 17, 1998, collection by D. Fowler and B. Mehl as part of the Warm Water Stream Survey. 1B shows location 11797 in May 2018, when the stream was not visible. 1C Shows the same location in April 2013. 1D Location 12985 (37.877713 N, 77.869453 W, also shown in April 2013) indicates four eels caught by Marcel Montane (VDGIF) on August 31, 1988. Location numbers are based on their row number in the database.

The land use raster dataset was cropped to the study area using the *Extract by Mask* function. Land use resolution was 300 m^2^. The raster originally contained twenty categories for land use, which were reduced to four for purposes of analysis using the *Reclassify* (Spatial Analyst) tool in ArcGIS. Areas marked “water bodies’ or “no data” (i.e. no land at that point) were reclassified as water. Areas marked “urban areas” were left as-is. Areas labeled “Cropland” and “Bare areas” were reclassified to one category called “cropland/barren”. All other categories, which included “Herbaceous cover”, “Mosaic cropland (>50%) / natural vegetation (tree, shrub, herbaceous cover) (<50%)”, “Mosaic natural vegetation (tree, shrub, herbaceous cover) (>50%) / cropland (<50%)”, “Tree cover, broadleaved, deciduous, closed to open (>15%)”, “Tree cover, broadleaved, deciduous, closed (>40%)”, “Tree cover, needle leaved, evergreen, closed to open (>15%)”, “Tree cover, needle leaved, deciduous, closed to open (>15%)”, “Tree cover, mixed leaf type (broadleaved and needle leaved)”, “Mosaic tree and shrub (>50%) / herbaceous cover (<50%)”, “Mosaic herbaceous cover (>50%) / tree and shrub (<50%)”, “Shrubland”, “Grassland”, “Sparse vegetation (tree, shrub, herbaceous cover) (<15%)”, “Tree cover, flooded, fresh or brackish water”, “Shrub or herbaceous cover, flooded, fresh/saline/brackish water” were reclassified as “forests, shrubs and mosaics”. While there are likely large differences between these areas, determining the effects of things like deciduous forest buffers vs. evergreen forest buffers on American Eel is beyond the scope of this work.

To analyze the variables of eel density, dam density and land use we created a ranking system from 0-3 for each variable. Eel and dam densities were calculated by their abundance in the subwatershed segments (43 in total) divided by the total area of each segment. Eel data were tabulated by adding up the segments upstream to downstream, on the assumption that if an eel was found upstream it would have had to swim through the downstream sections to reach that location. So, the eel density numbers for the lowest reach of the Potomac River includes all sections above it. Eel density data (km^-2^) were classified into four categories ranked lowest to highest: <1.0, ≥1 to 2, >2 to 5 and >5. Dams were calculated in the opposite order, downstream to upstream, because to get to the upper reaches of a river an eel would have to pass through the downstream sections first. The methodology to sum eel and dam densities (going from upstream to downstream and downstream to upstream) is illustrated in Fig. 2. Dam densities (100 km^-2^) were classified by subdividing the datasets to get a more or less even distribution of each of the four categories ranked highest to lowest: < 1, ≥1 to < 2, 2 to < 5 and ≥ 5 (Table 1). The limitation to this method is that many of the dams are not on the mainstems of the rivers so an eel traveling upstream could swim past them, but it provides a baseline of the difficulty of each section.

**Fig. 2.**
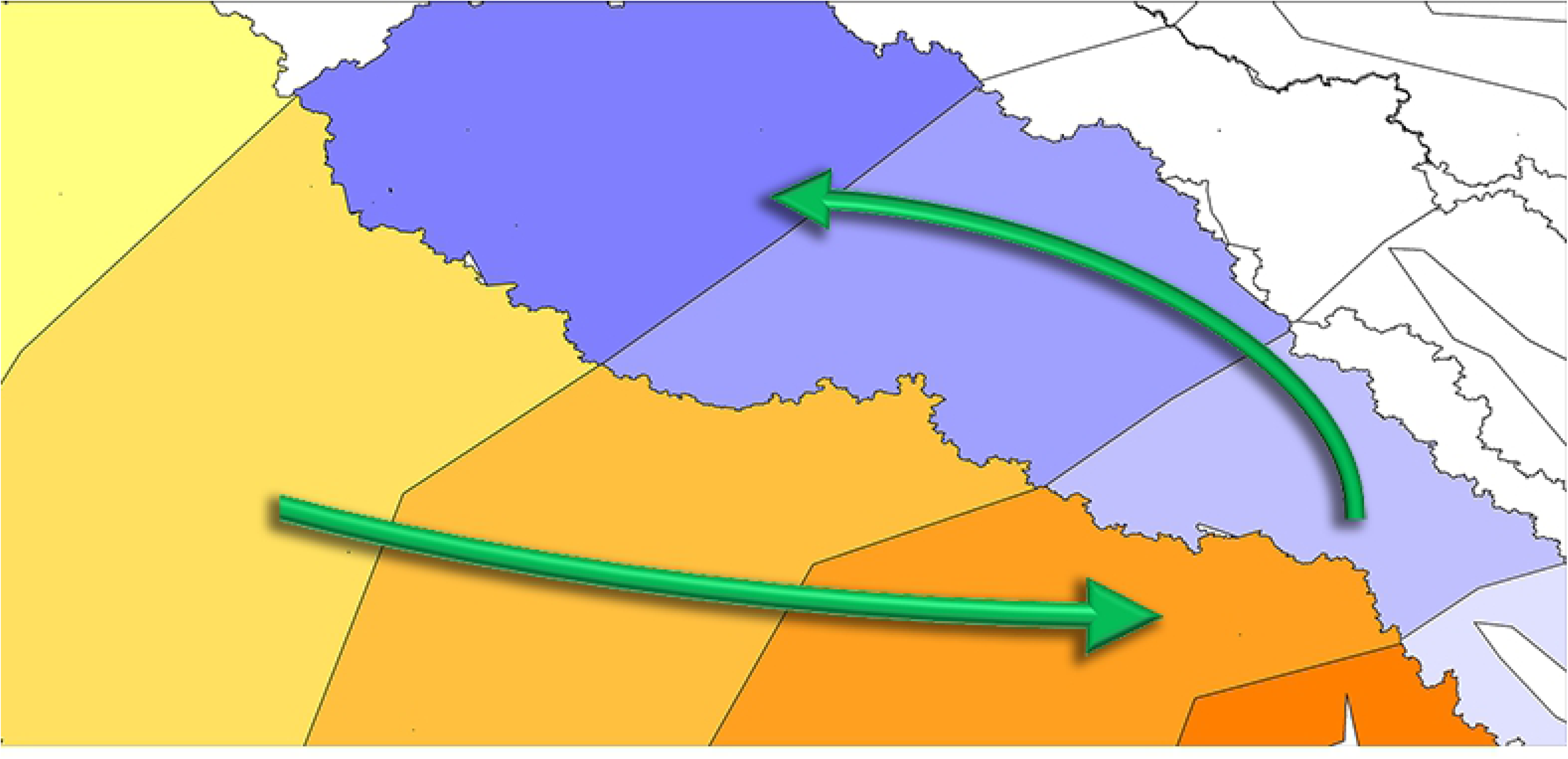
The methodology for summing data along watersheds. Eel density data are summed upstream to down (orange). Dam density data are summed downstream to up (lavender/blue).

**Table 1.** Categories for each variable and their assigned rankings

| **Ranking** | **0** | **1** | **2** | **3** |
| --- | --- | --- | --- | --- |
| Eel density (km<sup>-2</sup>) | < 1.0 | ≥ 1.0 to <2.0 | ≥2 to 5 | > 5 |
| Dams w/o fish passage (100 km<sup>-2</sup>) | ≥ 5 | 2 to < 5 | 1 to < 2 | < 1 |
| Land use | Urban areas | Cropland/barren | Forest, shrubs and mosaics | Water |

The subwatershed polygon colors were adjusted using the *Symbology* tab in ArcGIS. By using the option to show *Unique Values* and a color ramp that runs from red to blue, new feature classes were generated from two value fields: eel density and dam density. These were converted from shapefiles to rasters using *Feature to Raster* (Conversion). This was not necessary for land use because it was already in raster format.

Each variable (eel density, dam density and land use) has four possible values (0-3), thus each raster has 4 colors. With two variables there are 16 possible values (4^2^) and with three variables there are and 64 (4^3^) possible values, which can be represented as 4-bit (2^4^) or 6-bit (2^6^) rasters. Simply summing the rasters together however will not produce this range of values, because the numbers will overlap and even combining all three layers will produce only 10 possible outcomes (0-9). Thus, the eel and dam density rasters were weighted using the *Reclassify* function to 1) account for the importance of each variable and 2) to increase the color depth of the output rasters.

Of the three variables, eel density was given the highest priority, because if eels were found then by definition the area is habitable by eels. (Density was used instead of CPSE for this section, because it allows a much larger dataset with more coverage of each subwatershed segment). Dams are second, because barriers that physically block fish passage have been found to have a greater impact on eel abundance and density than urbanization [14]. Eel density was reclassified by multiplying the values by 16 (2^4^) and dam abundance was reclassified by a factor of 4 (2^2^). This should not be interpreted as dams having exactly four times the importance of land use and eel abundance having four times the importance of dams. These numbers are used so that each combination of variables produces a unique value with no overlaps, thus maximizing the color depth of the raster when combining variables (in the case of three variables, it results in more than a six-fold increase). All of the possible values are represented in a binary matrix (Table S1). To the best of our knowledge, this is the first study to sum multiple 2-bit rasters as binary numbers to generate a color palette for visual representation of data. Rasters were summed together using the *Cell Statistics* (Spatial Analyst) function in ArcGIS.

CPSE was calculated separately and not included in the preceding analysis. To calculate CPSE within the study area, we filtered the eel data by removing any fish identified as glass eels or elvers (which are much smaller and thus likely to be caught in higher numbers than yellow or silver eels) and removed any counts >99 at a given sample point, because these may have also represented unlabeled glass eels/elvers. This created a subset of 70,794 eels, primarily between the years 1977 and 2015, which we then identified by subwatershed by using the Clip (Analysis) tool in ArcGIS. The totals in each subwatershed were divided by the number of sampling events and the result (including zero counts if applicable) was considered the CPSE. CPSE was compared to eel and dam density using scatterplots and linear regression.

## Results

Based on the Digital Elevation Model data, we created 24 subwatersheds (Table S2). The watersheds labeled by their ID numbers are shown (Fig. 3).

**Fig. 3.**
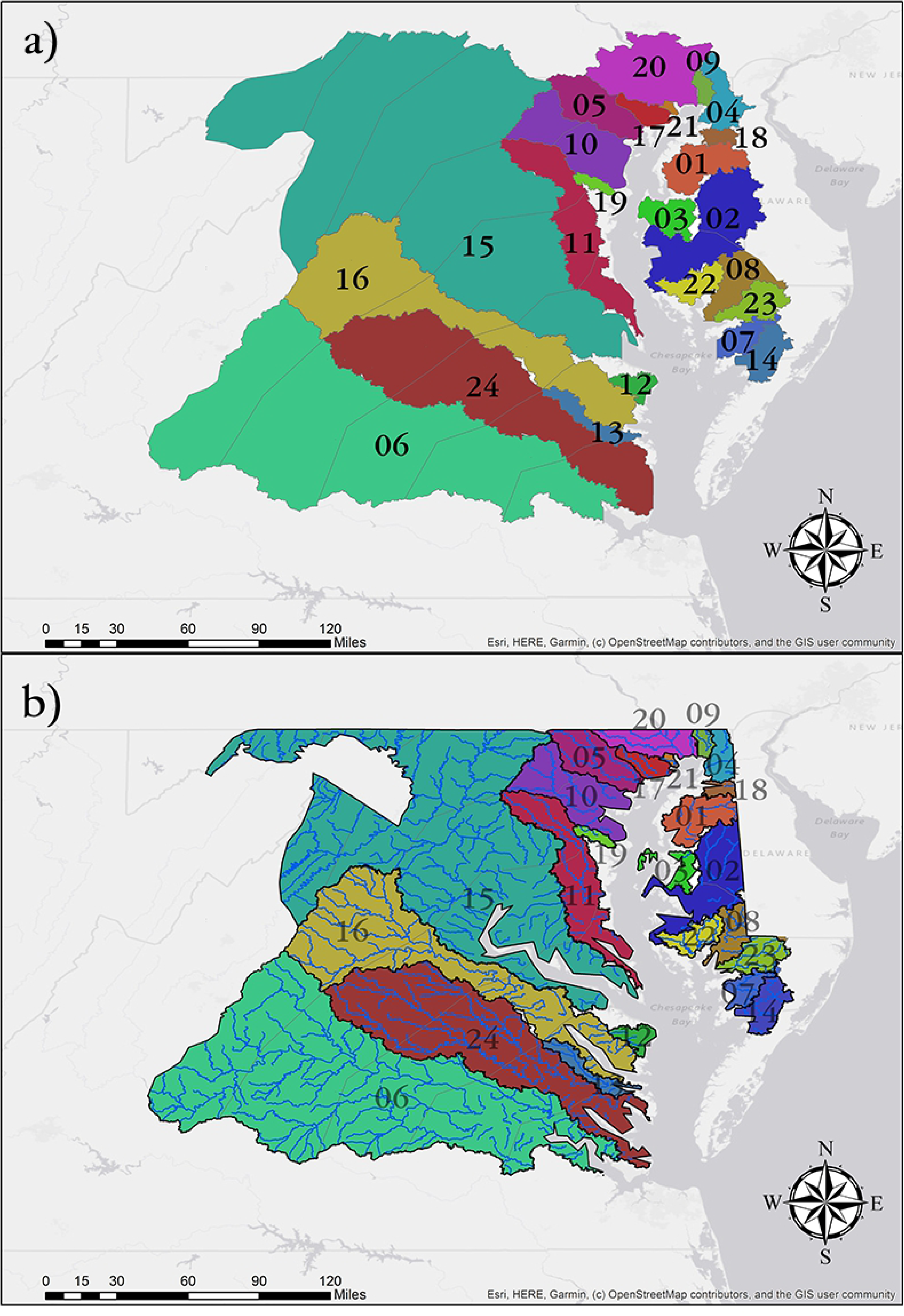
Chesapeake Bay subwatersheds generated from a Digital Elevation Model (DEM). 3a shows the watersheds using the same ID numbers from Table S2 as labels: 01 – Chester River, 02 – Choptank River, 03 – East Bay, 04 – Elk River, 05 – Gunpowder River, 06 – James River, 07 – Manokin River, 08 – Nanticoke River, 09 – North East River, 10 – Patapsco River, 11 – Patuxent River, 12 – Penny Creek, 13 – Piankatank River, 14 – Pocomoke River, 15 – Potomac River, 16 – Rappahannock River, 17 – Romney Creek, 18 – Sassafras River, 19 – Severn River, 20 – Susquehanna River, 21 – Swan Creek, 22 – Transquaking River, 23 – Wicomico River and 24 –York River. 3b shows the watersheds after being clipped to the boundaries of Virginia, Maryland and Washington, D.C., with rivers based on the DEM data. Not all rivers and tributaries are shown.

**Table 2.**
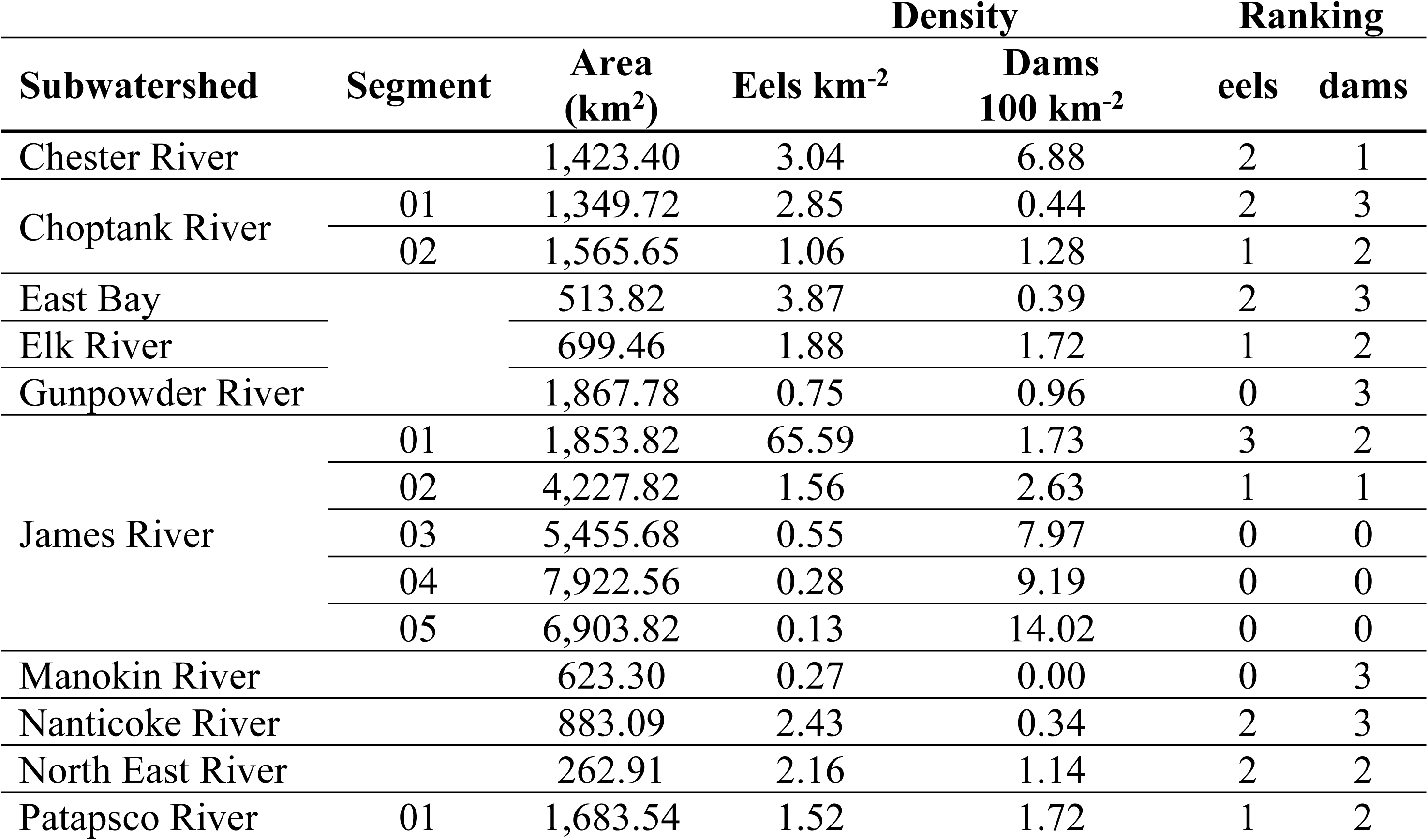
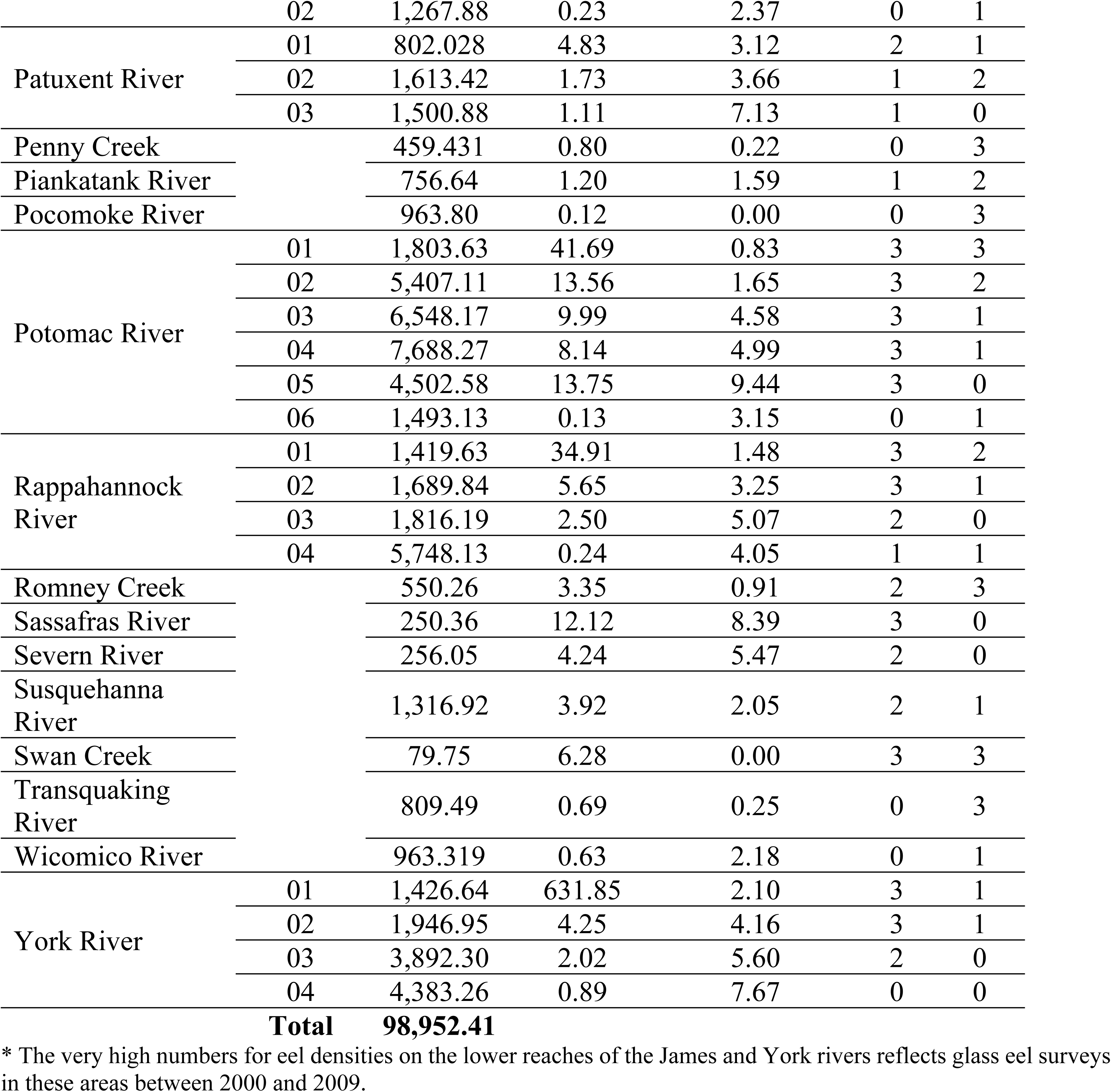
Area (km^2^), eel density (km^-2^), dam density (100 km^-2^) and rankings for each subwatershed segment

Delineating the watersheds at approximately 50 km along the mainstem resulted in multiple segments for six of the rivers and 43 segments in total. Segment 01 starts at the mouth of the river where it enters the Chesapeake Bay and the numbers increment from downstream to upstream (Table 2).

There were 1,184,141 eels sampled in the study area. These data were clipped to the study area (Fig. 4).

**Fig. 4.**
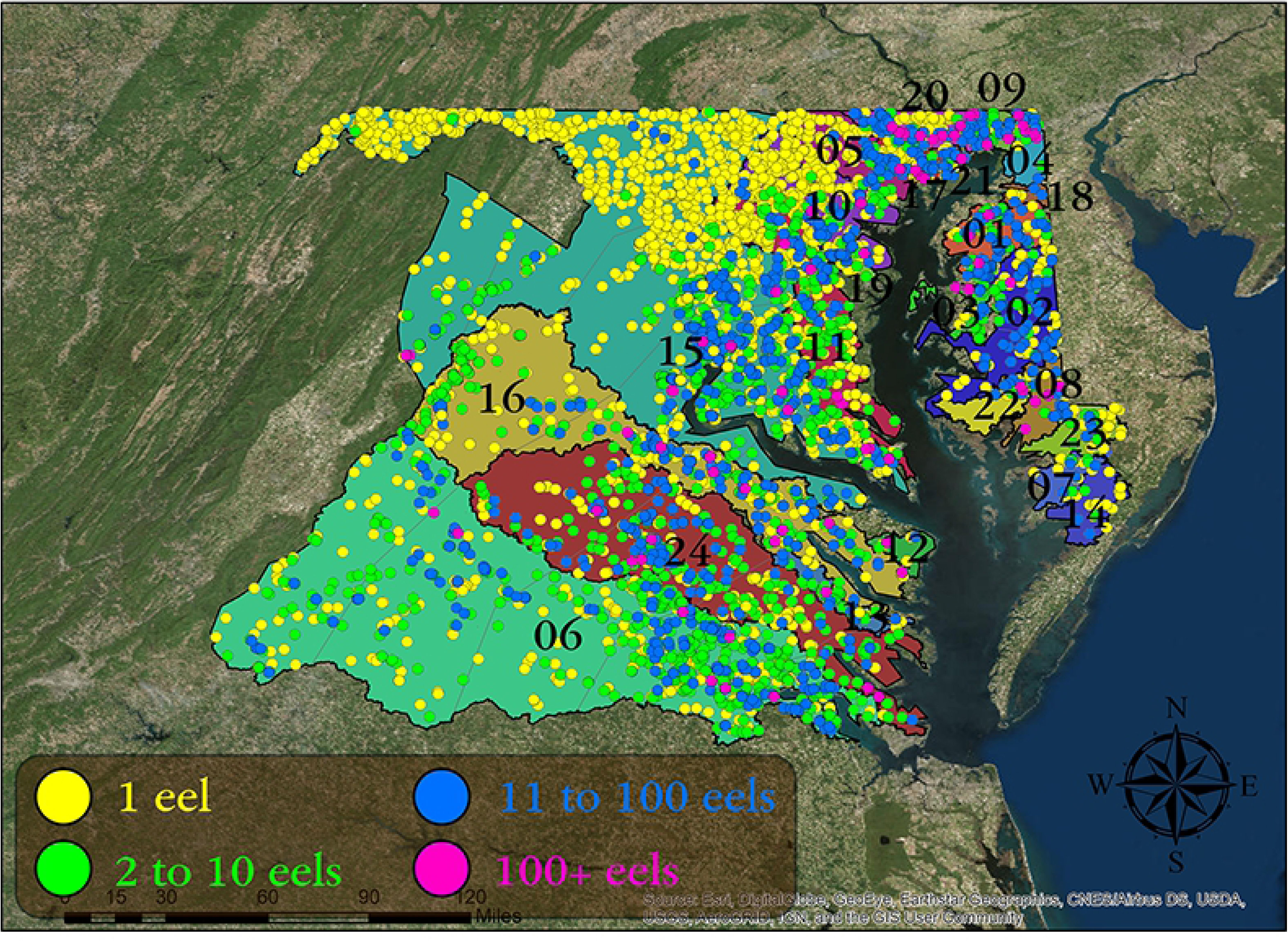
American Eel (*Anguilla rostrata*) abundance data clipped to the study area of Chesapeake Bay subwatersheds. Yellow circles represent individual eels, green 2 to 10 eels, blue 11 to 100 and magenta 100 or more. Numbers correspond to watershed labels in Figure 3.

The study area had 2,435 dams, with 30 (1.23%) having provisions for fish passage (Fig. 5). The date of construction was available for 913 of the dams. In the study area, all dams were built between 1800 and 2001 (Fig. 6) and the highest years of dam construction were from 1955 to 1975, with an average of 26 dams built per year during these two decades (Fig. 7). The graph does not include records of any dams that were removed.

**Fig. 5.**
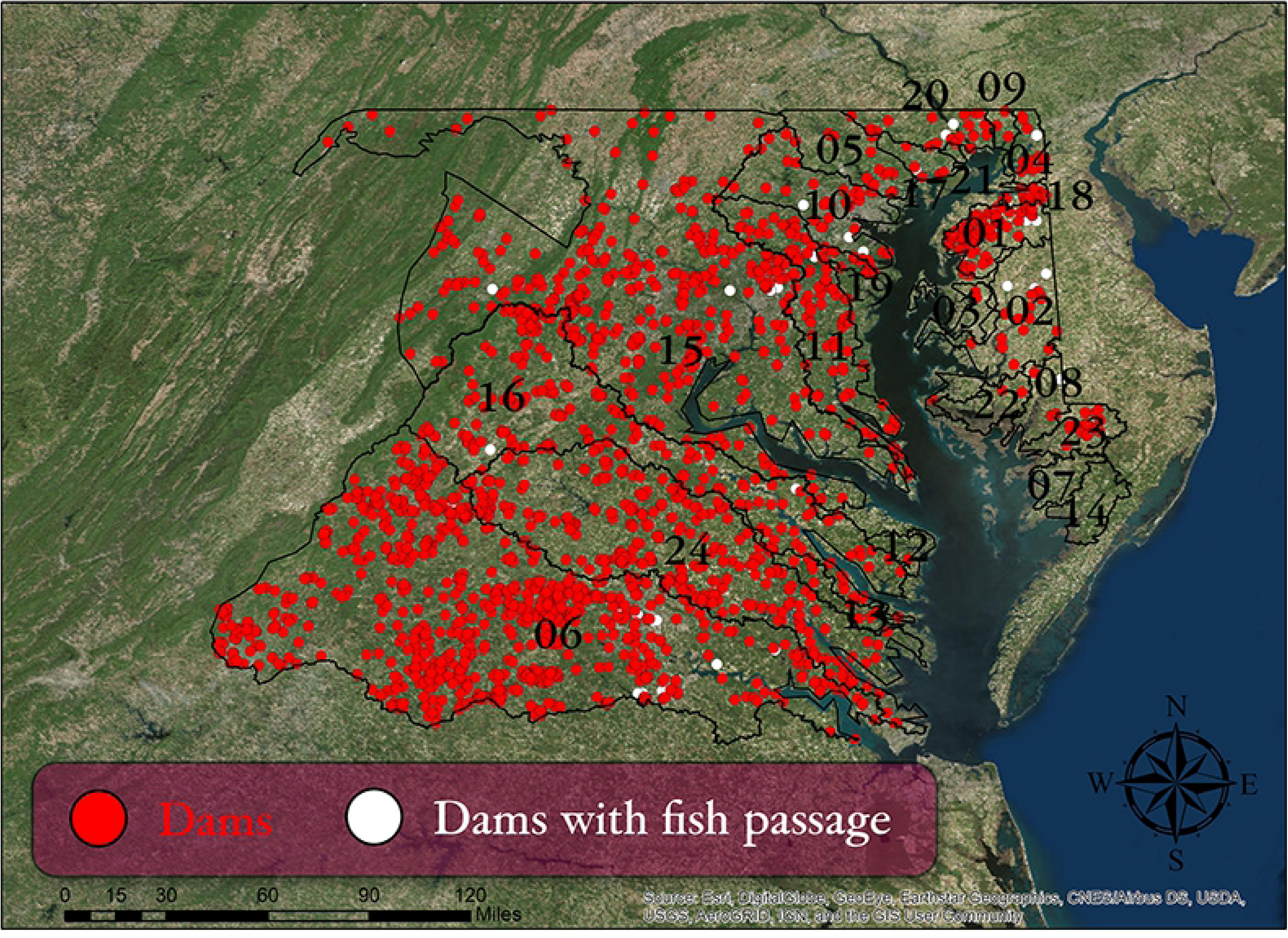
Locations of dams in the study area (Chesapeake Bay subwatersheds). Red dots indicate dams without fish passage; white dots indicate dams with upstream fish passage. Numbers correspond to the watershed labels in Figure 3.

**Fig. 6.**
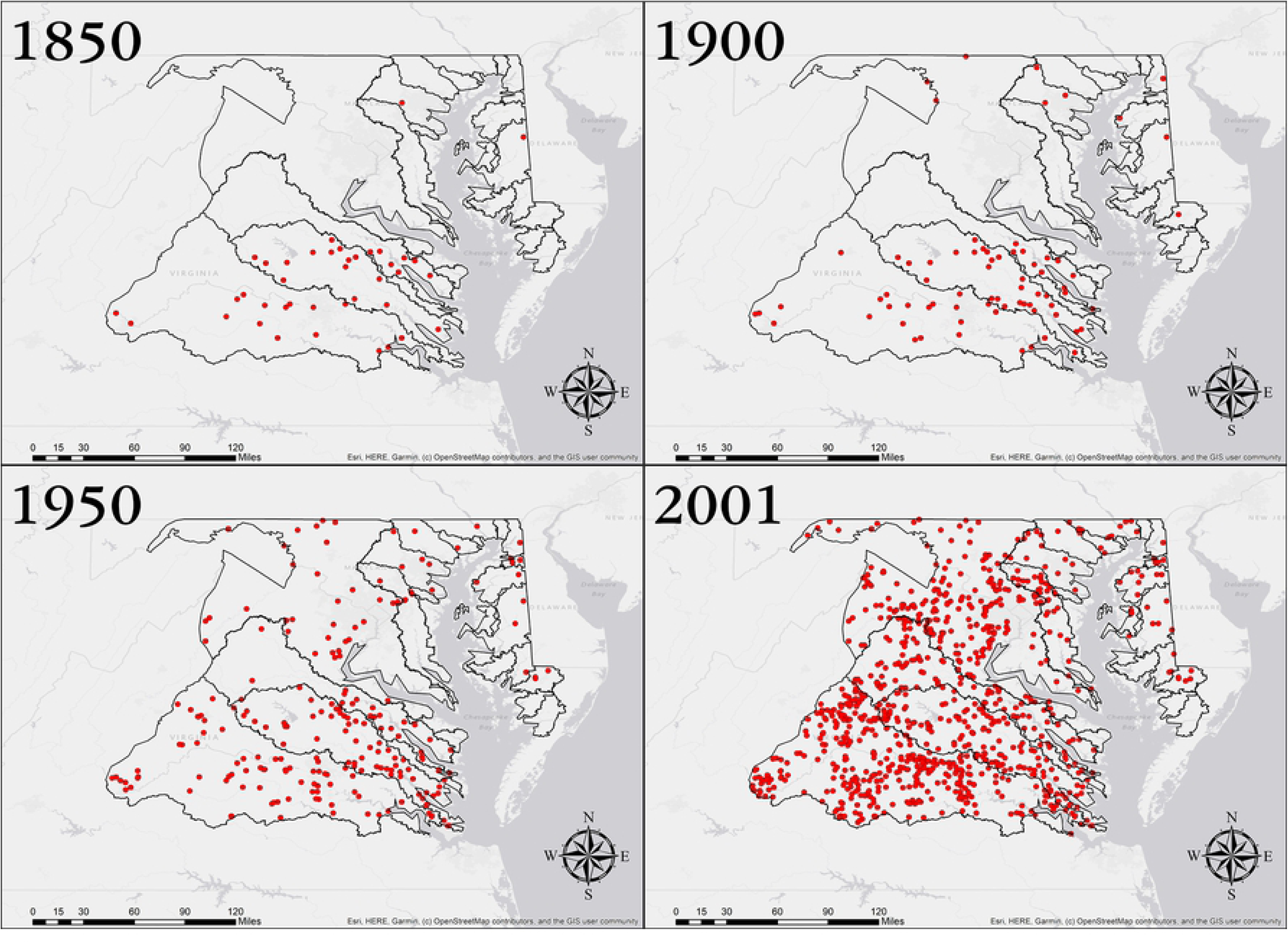
Dams over time in the study area (Chesapeake Bay subwatersheds) for the 913 dams where date of construction was available.

**Fig. 7.**
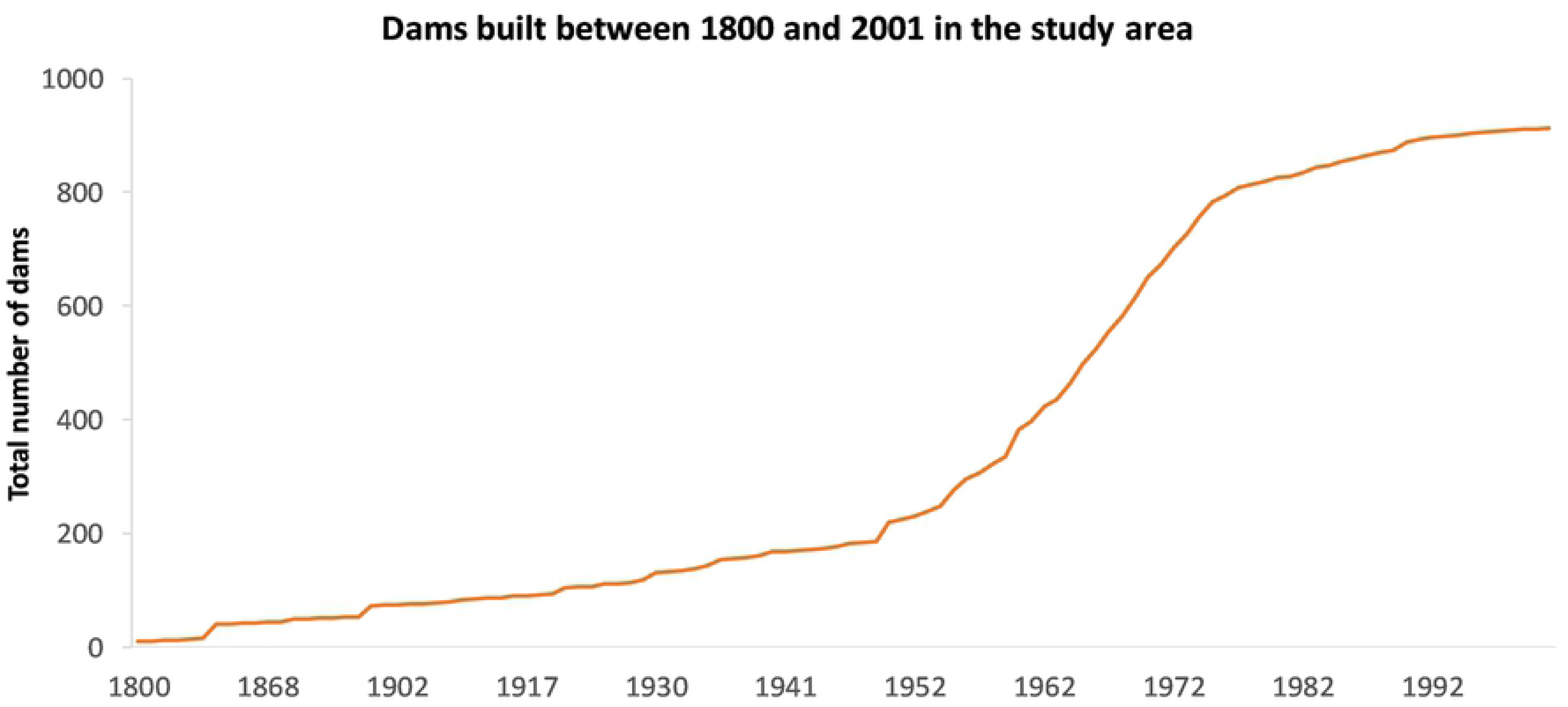
Construction of dams in the study area (Chesapeake Bay subwatersheds) from 1800 to 2001 for the 913 dams where the date constructed was known. Note years of peak construction between 1955-1975.

Eel densities were highest in York River 02, James River 01, Potomac River 01-05, Rappahannock River 01-02 and the Sassafras River. They were lowest in the Gunpowder River, James River 03-05, Manokin River, Penny Creek, Pocomoke River, Potomac River 06, Transquaking River, Wicomico River and York River 04. Dam density was highest in the Chester River, James River 03-05, Patuxent River 03, Potomac River 05, Rappahannock River 03, Sassafras River, Severn River and York River 03-04. Dam density was lowest in the Choptank River, Gunpowder River, Manokin River, Nanticoke River, Penny Creek, Pocomoke River, Romney Creek, Swan Creek and the Transquaking River. Three of these locations (Manokin River, Pocomoke River and Swan Creek) have no dams at all (Fig 8.)

**Fig. 8.**
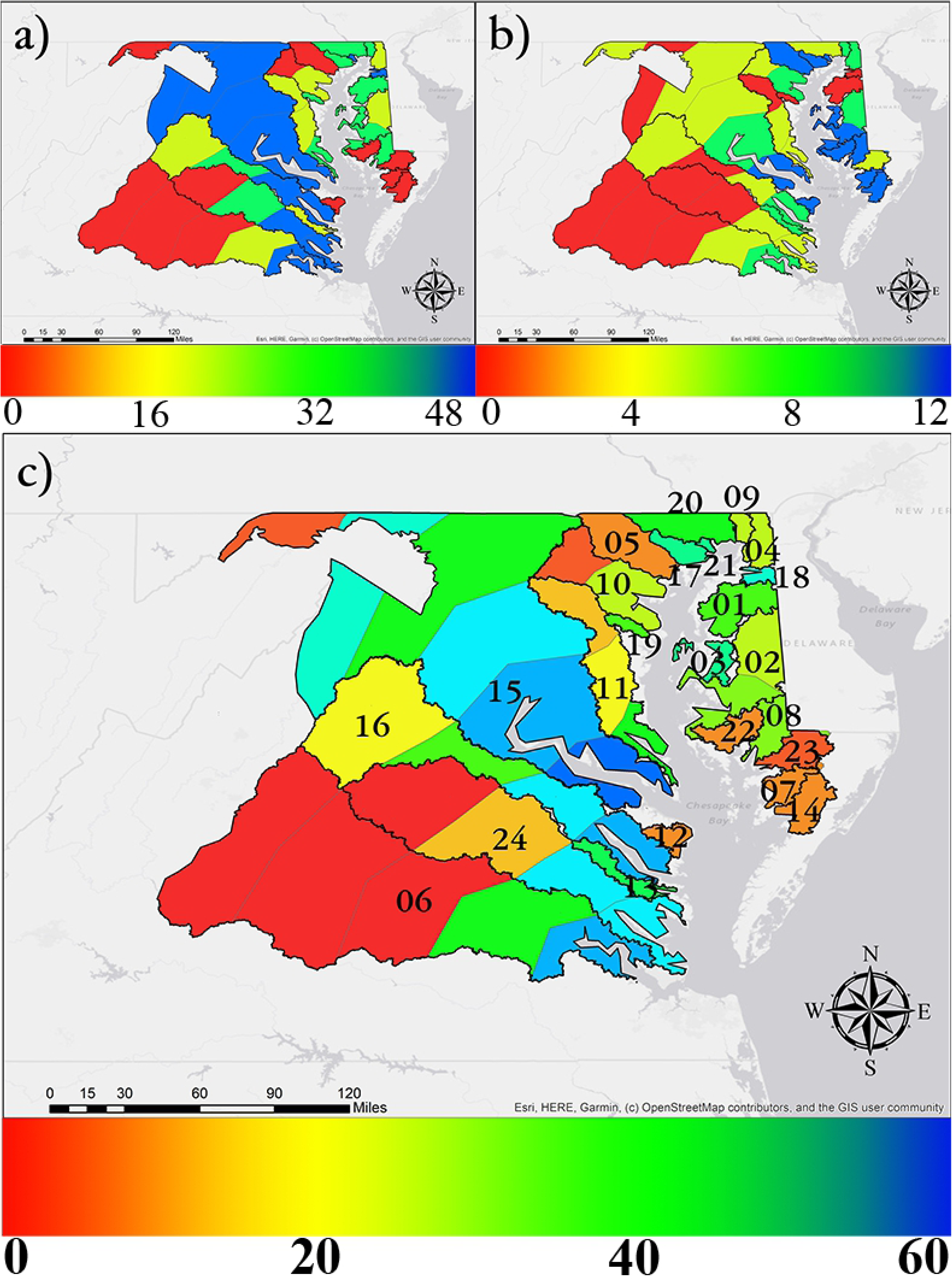
Rankings by eel and dam densities. 8a shows eel density, 8b shows dam density and 8c and shows the combined dam and eel density rankings. Color ramp indicates rankings from low (red) to high (blue). As shown in Table S1, potential values range from 0 to 60. Numbers correspond to watershed labels in Figure 3.

The reclassified land use data showed most (88.96%) of the land within 2 km of the rivers was classified as Category 2 – forests, shrubs and mosaics, followed by urban areas (5.44%), water (4.87%) and cropland/barren (0.73%) (Fig. 9).

**Fig. 9.**
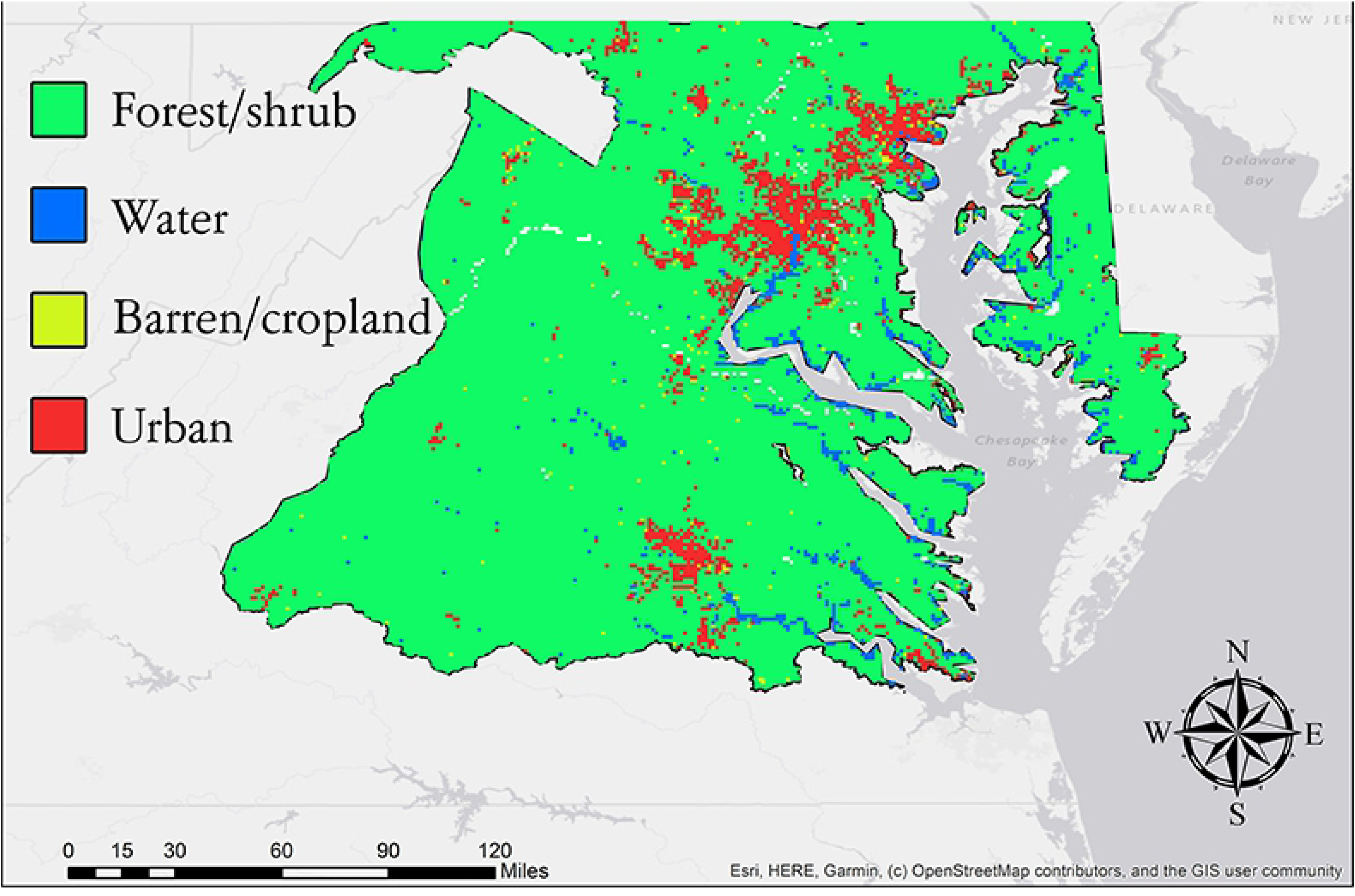
Land use categories. Green indicates areas with plant cover, blue represents water, yellow barren surfaces/cropland only and red urbanized areas. Red primarily corresponds to the three metropolitan areas on the map, Baltimore Maryland, Washington D.C. and Richmond Virginia.

Combining the rasters for dams and land use produces a 4-bit raster with 16 possible values for each pixel. Areas in blue indicate few barriers and good land use values while areas in red indicate areas with more barriers and impervious surfaces or barren areas (Fig. 10).

**Fig. 10.**
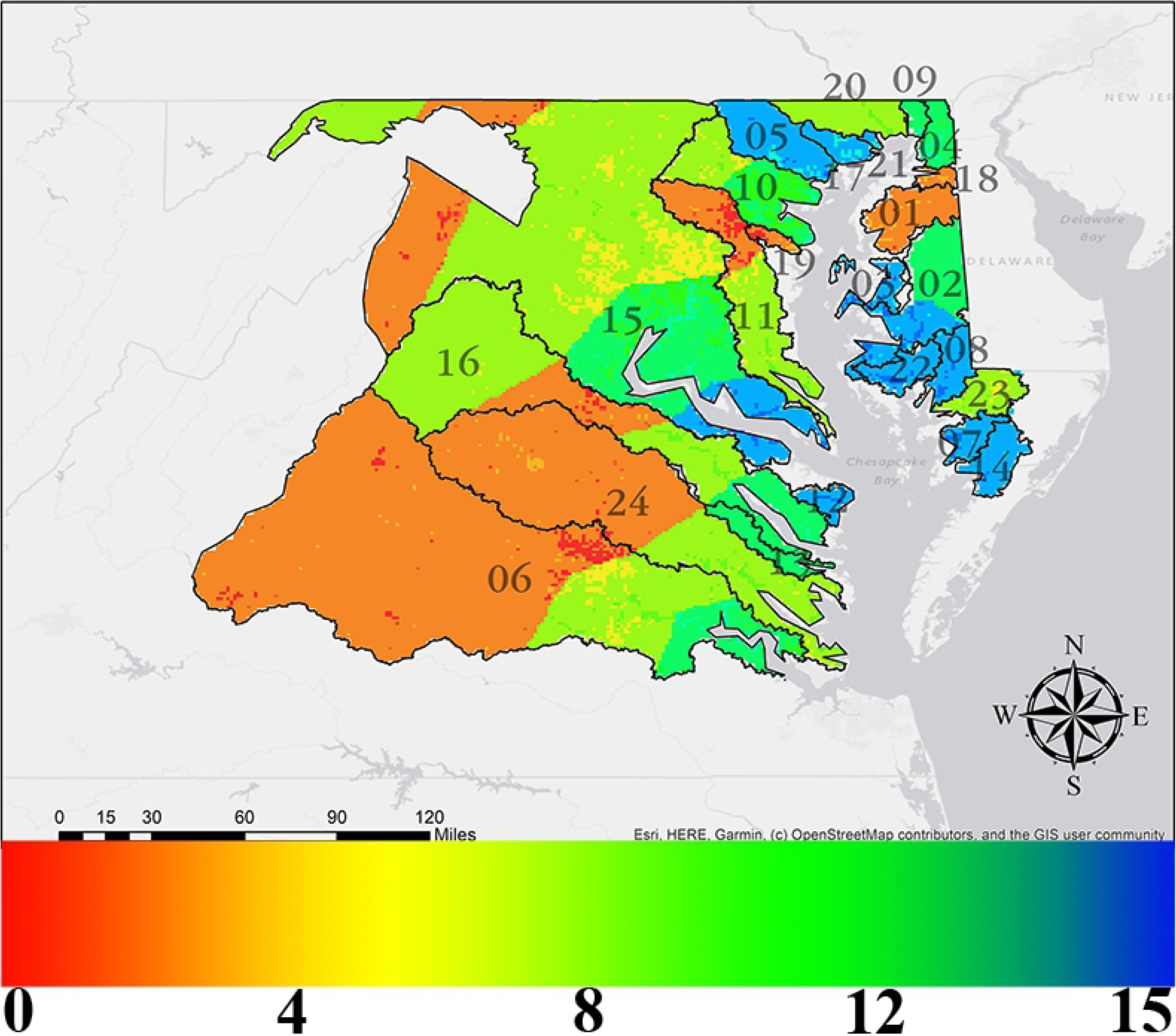
A “features only” map combining the rankings for dam density and land use. The scale runs from blue (highest-ranked areas with fewer dams and higher ranking land use categories) to red (lowest-ranked areas with more dams and lower ranking land use categories). As shown in Table S1, potential values range from 0 to 15. Numbers correspond to watershed labels in Figure 3.

Stream boundaries were generated from the DEM (dark blue lines) and combined with the USGS data, with a 2 km polygon around the polylines (Fig. 11).

**Fig. 11.**
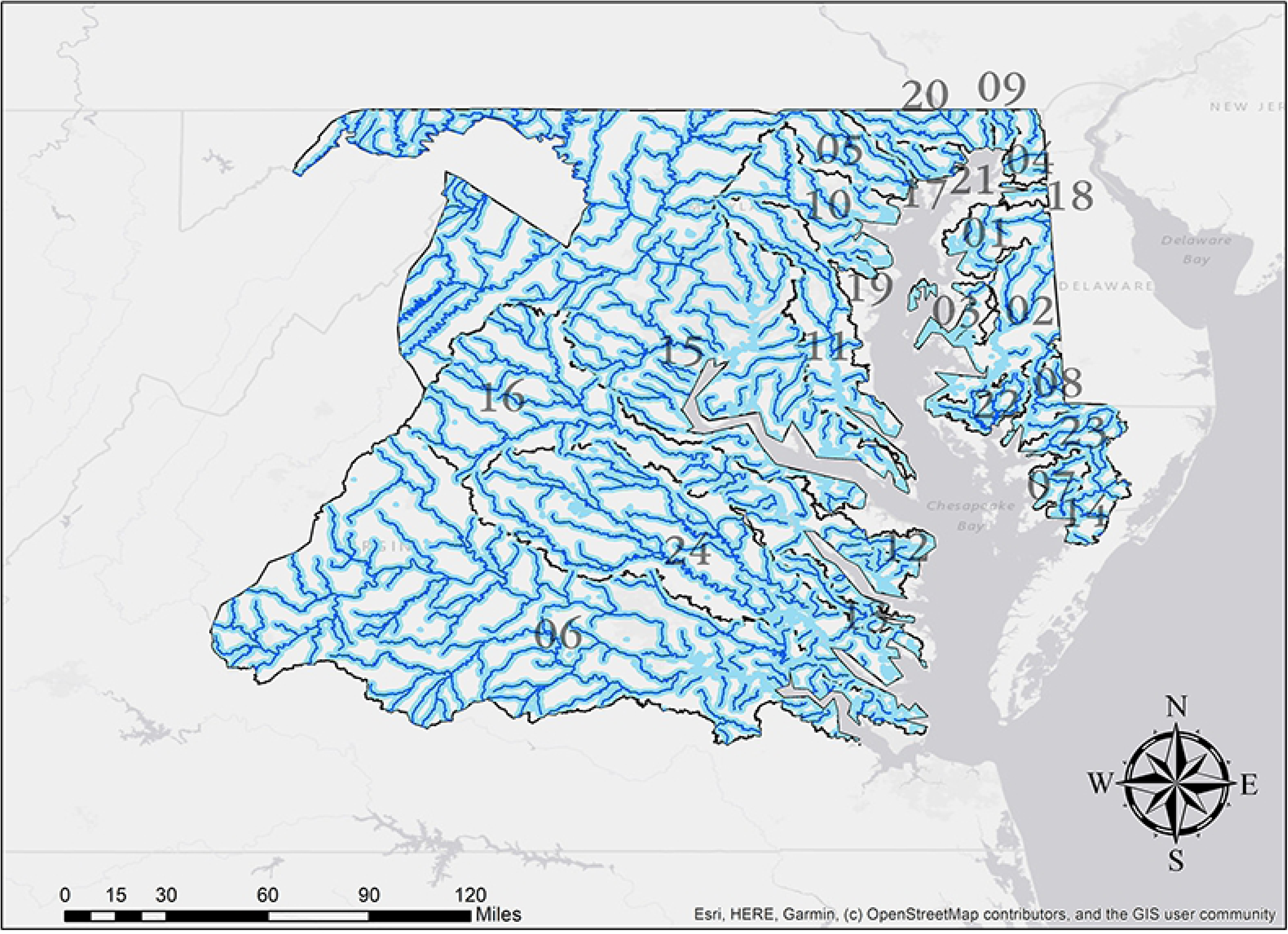
Streams clipped to study area boundaries. The darker blue lines are generated from the Digital Elevation Model (DEM), while the lighter blue lines are a combination of the USGS imagery, the DEM and a 2 km polygon around the polylines. Numbers correspond to watershed labels in Figure 3.

When the three layers (eel density, dam density and land use) are added together, the result is a 6-bit (64 possible colors) raster. This raster was extracted from the stream boundary mask to produce Fig. 12.

**Fig. 12.**
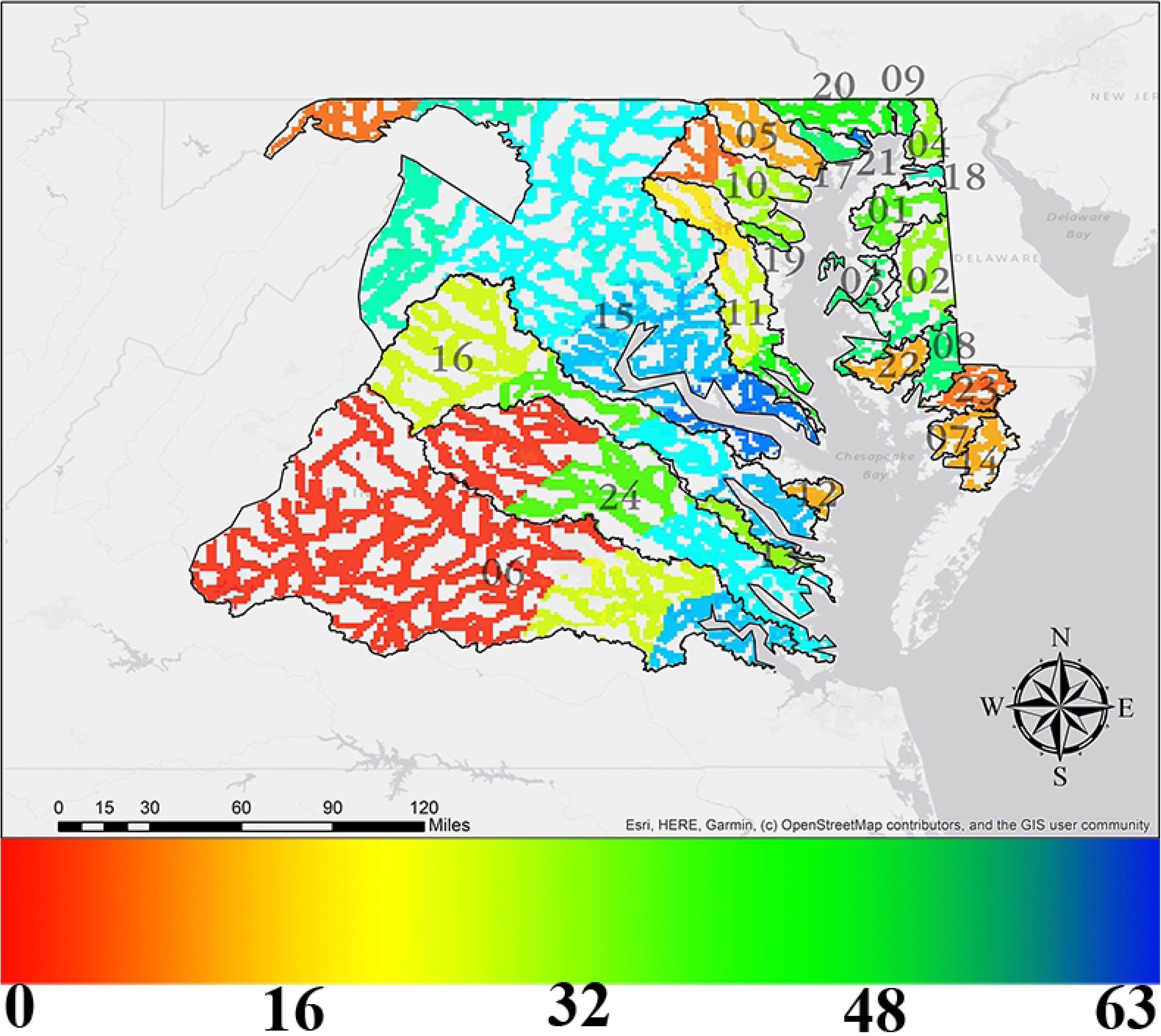
Map combining the rankings for eel density, dam density and land use clipped to stream boundaries. The color bar runs from red (lowest-ranked areas) to blue (highest-ranked areas) with values from 0 to 63. Numbers correspond to watershed labels in Figure 3.

CPSE varied by watershed and segment. The average CPSE was 7.78, with a standard deviation of 4.88. The lowest value was 1.05, at Potomac River 06 and the highest value was 23.38 at York River 01 (Fig. 13). CPSE was compared to the density of yellow/silver stage eels (Fig. 14). The coefficient of determination (R^2^) is 0.657, thus the coefficient of correlation (R) is 0.810. CPSE was also compared to the density of dams (Fig. 15) per subwatershed segment.

**Fig. 13.**
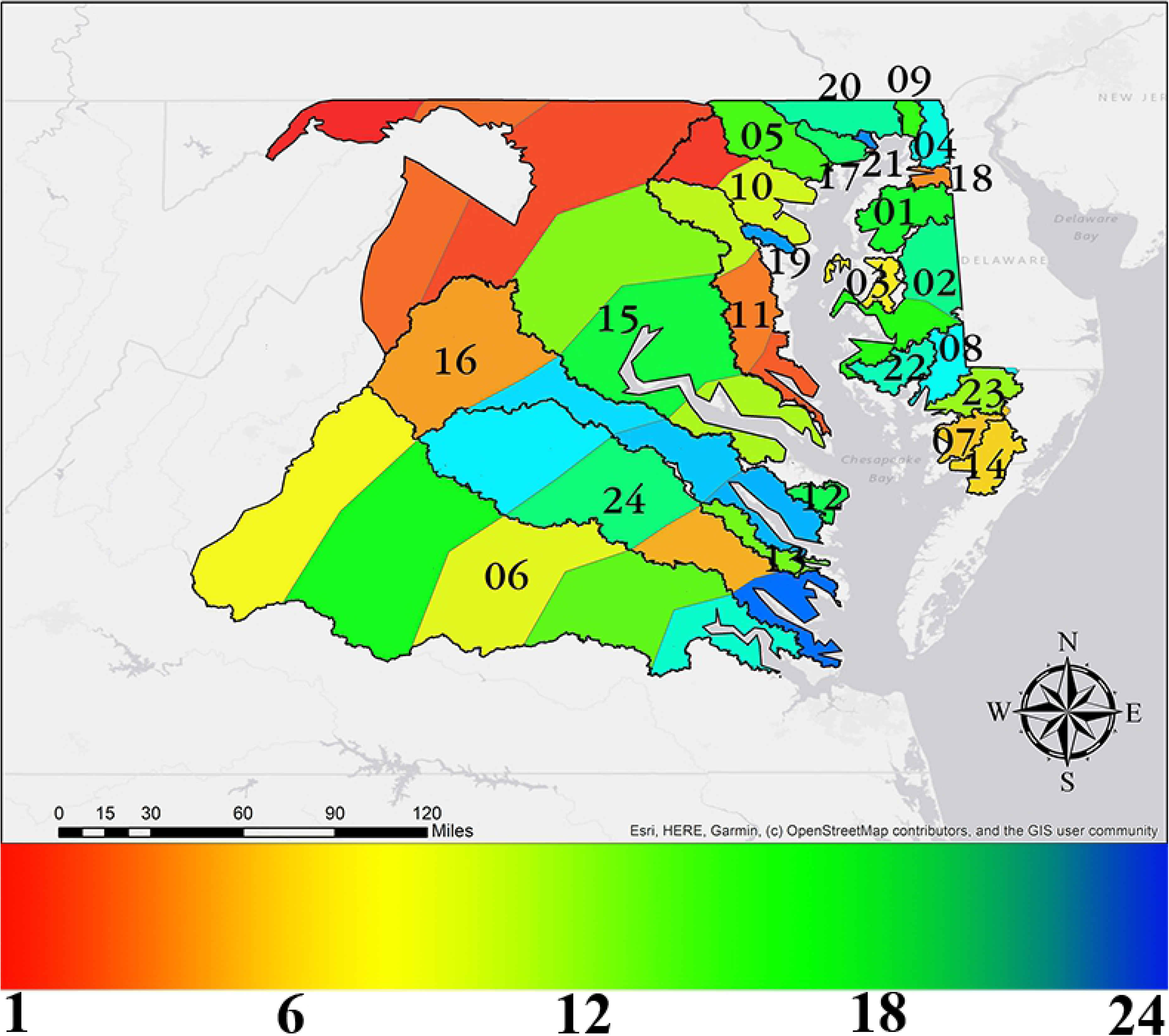
Catch per sampling event (CPSE) for all watershed segments. Values range from red (low) to blue (high). Colors correspond to the CPSE values themselves rather than rankings. Numbers correspond to watershed labels in Figure 3.

**Fig. 14.**
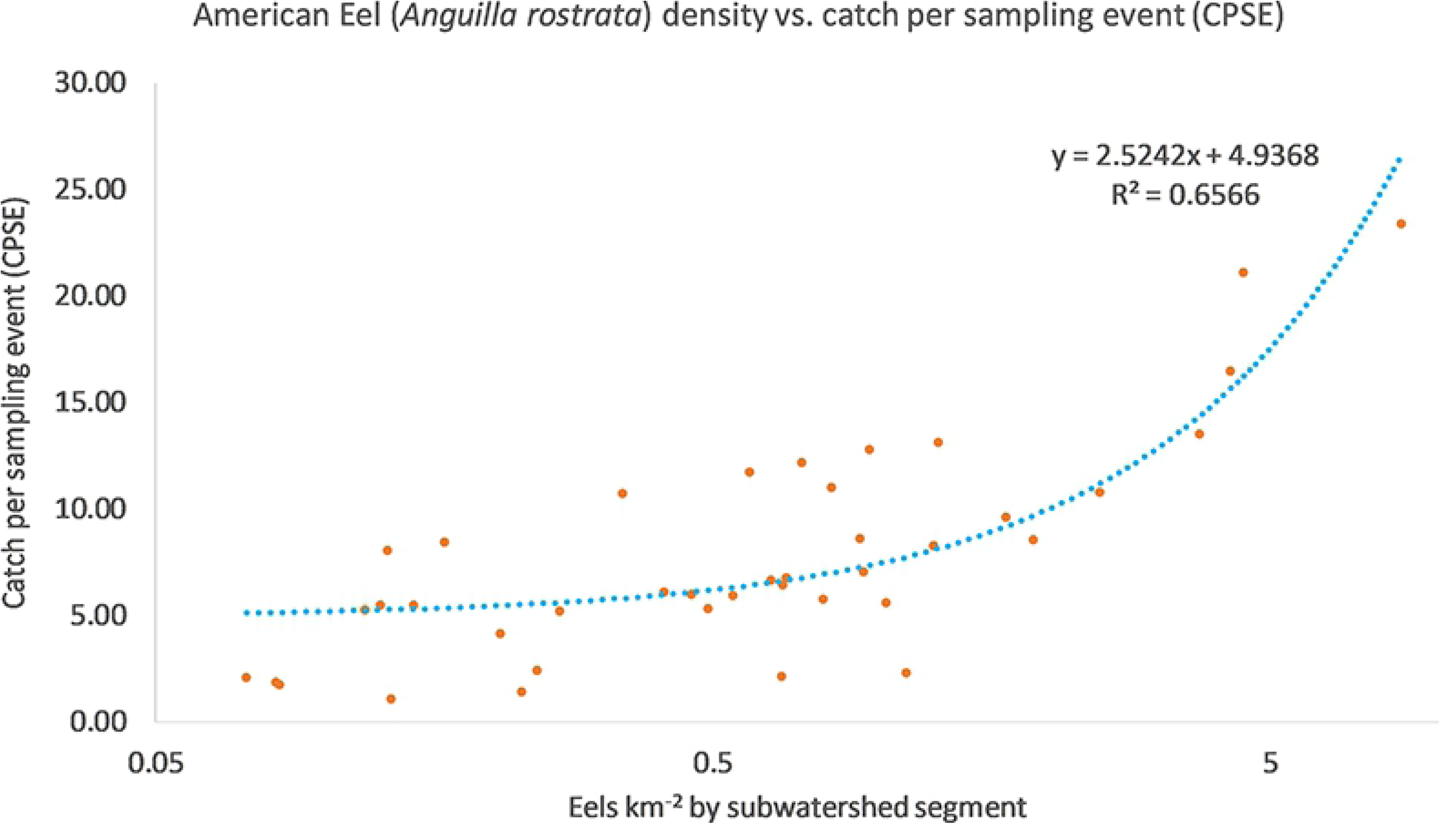
Yellow and silver stage American Eel (*Anguilla rostrata*) density vs. Catch per sampling event (CPSE) by subwatershed segments. The dot in the upper right corner is York River 01, which had the highest eel density (8.62) and the highest CPSE (23.38). Trendline is based on linear regression (note that the line is curved because the x-axis is log scale).

**Fig. 15.**
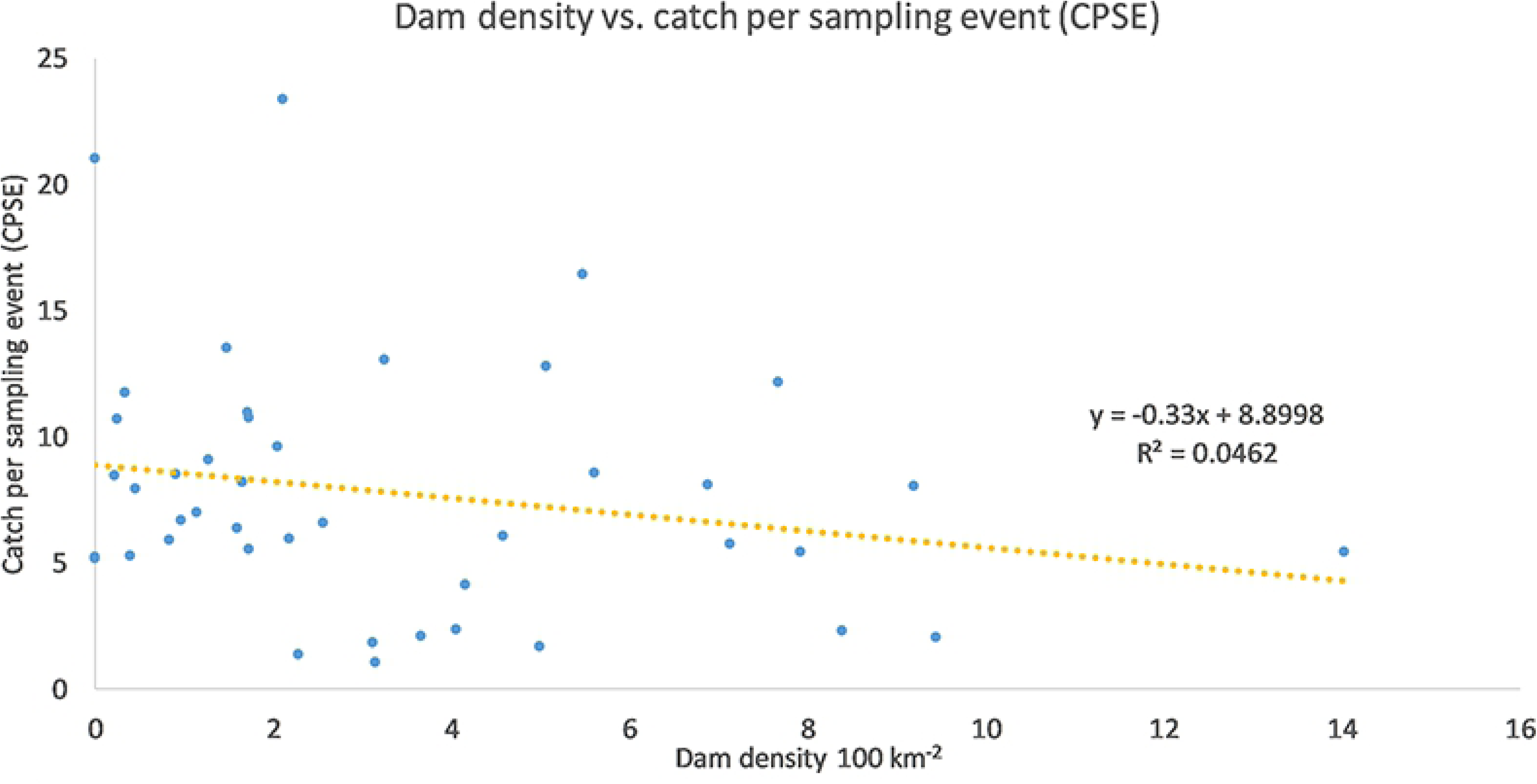
Dam density (100 km^-2^) and catch per sampling event per watershed. Trendline is based on linear regression.

## Discussion

As demonstrated by these findings, American Eels have been sampled in all major drainages of the Chesapeake Bay, although density and CPSE vary greatly by subwatershed segments. Dams were mapped and their impacts can be assessed not only locally but across entire rivers. Our model also includes the localized effects of land use. The primary contribution of this work is the first creation of a toolset for spatial analysis applicable to eels across a wide geographic range and (with minor modifications) other diadromous fishes. The benefit of using spatial analysis in species conservation is to be able to visualize trends over space and time. This will be essential in determining what caused the decline in the eel population; if it was primarily due to fishing, we might expect to see similar declines everywhere, on the other hand if local factors such as dams and land use are more important factors then the declines should be more pronounced in those areas.

There were some limitations to this study including that the eel data collection records were not consistent by region and CPSE presents its own limitations. Thus, based on results we cannot definitively say if one river or segment has a higher eel density than another and areas in red should be considered candidates for further study rather than places where eels are disappearing. The eel numbers are baseline records of what has been documented, thus a river that appears to have low eel density could potentially improve with more consistent sample methods. On the other hand, areas with relatively high densities can be considered as places where eels appear to be doing well. Additionally, the dam density and land use data can be used to determine eel habitat suitability.

One benefit to dividing the rivers into segments and adding eel abundance cumulatively to create the total eel densities is seen on the Potomac River, where the downstream locations do not have many documented eels but the numbers from the Shenandoah tributary (Segment 05) make up for it. Thus, this methodology goes beyond tallying eel counts in a localized area and shows how sampling in one place can affect the ranking of an area 200 km away. This has implications for restoring watersheds, because removing a dam downstream may have impacts felt over a large geographic area. This may be especially important for cost/benefit analyses of constructing dams or fish passages.

Based on the eel density layer only, the highest ranking (> 5 eels km^-2^) subwatershed segments were James River 01, Potomac River 01-05, Rappahannock River 01-02, Sassafras River, Swan Creek and York River 01-02. The lowest ranking (< 1.0 eels km^-2^) areas were Gunpowder River, James River 03-05, Manokin River, Patapsco River 02, Penny Creek, Pocomoke River, Potomac River 06, Transquaking River, Wicomico River and York River 04. Two locations (James River 01 and York River 01) have very high eel densities because of glass eel surveys in these areas between 2000 and 2009. This however does not impair the accuracy of the results because these locations would have had the maximum value for eels (3) even without the glass eel counts.

Examining only the dam density layer, the highest ranked areas (dam density < 1 100 km^-2^) were Choptank River 01, East Bay, Gunpowder River, Manokin River, Nanticoke River, Penny Creek, Pocomoke River, Potomac River 01-02, Romney Creek, Swan Creek and Wicomico River. Three of these locations (Manokin River, Pocomoke River and Swan Creek) had no dams in the study area. The areas with the highest dam densities (≥5 100 km^-2^) were the Chester River, James River 03-05, Patuxent River 03, Potomac River 05, Rappahannock River 04, Sassafras River, Severn River and York River 03-04.

When the eel and dam data were combined, the areas of most concern were York River 04 (upstream of Cedar Fork, VA) and James River 03-05 (from the headwaters to Richmond), especially Segment 05, which had the highest dam density (14.02 100 km^-2^). Each of these areas had the lowest eel density rankings and the highest dam density rankings. Wicomico River and Patapsco River 02 also had the lowest rank for eel densities and second highest dam density ranking. Other areas appeared to be doing comparatively better. Potomac River 01 and Swan Creek each had the highest eel density ranking and the lowest dam density ranking. Swan Creek also had the second highest CPSE, 21.06, and was the only location besides York River 01 to have a CPSE >20. Potomac River 02, James River 01 and Rappahannock River 01 each had the highest eel density ranking and the second lowest dam density ranking. Potomac River 03, Rappahannock River 02 and York River 01-02 had the highest eel density ranking and the third lowest dam density ranking.

When eel density and CPSE were compared by watershed segments (Fig. 14), there was a positive relationship based on the correlation coefficient (R = 0.810). This could indicate the eel density data accurately reflects actual eel abundance, but without ground truthing in the form of sampling eels in these areas using consistent gear and methodology, it is difficult to draw further conclusions from these data. There may be a slight negative relationship between dam density and CPSE (Fig. 15) but further studies will be necessary to make this determination.

Based on these results, the Potomac River and Rappahannock River appear to be watersheds of lesser concern, although the Potomac River drops to a very low ranking at the upper reach (Segment 06). This could be due to reduced sampling effort above Segment 05, an area that was sampled many times by the USFWS, or it could be the result of the dams in Segment 05 (9.44 100 km^-2^). Yet another complication is that this section of the Potomac was missing large portions of the watershed in PA and WV, which are outside the study area because we did not look at eel data from these states. It would be beneficial to run this analysis including eel data from all Chesapeake Bay states and the full extent of the Potomac River before drawing conclusions on habitat suitability of the upper regions of this river. The James River and York River both start with good rankings for eel and dam densities near the Bay but end up with much lower scores further upstream, indicating barriers may be affecting upstream eel densities.

The rankings for eels and dams did not correlate at most locations and are sometimes contradictory. For example, Gunpowder River, Manokin River, Penny Creek, Pocomoke River and Transquaking River all had very low eel densities despite also having low densities of dams. In these cases, the low eel densities could reflect a lack of sampling efforts in these locations, or that there are reasons other than dams for the low numbers. The latter explanation seems more likely for the Gunpowder River, Manokin River and Pocomoke River, since each location had below average CPSE, whereas Penny Creek and Transquaking River had above average CPSE.

On the other hand, Potomac River 05 and Sassafras River have very high eel densities while also having high dam densities, indicating that at least in some cases the dams are not acting as complete barriers to migration, which is supported by previous studies [9]. This is especially interesting considering both locations have below average CPSE.

Land use was highly localized, with the majority of the urban areas (impervious surfaces) occurring in the metropolitan areas around Baltimore MD, Washington D.C. and Richmond, VA. These correspond to the subwatersheds of James River 02-03, Potomac River 02-03, Patuxent River 02-03, Severn River and Patapsco River 01.

Eel data was weighted more heavily than either of the other factors, because while stream connectivity and riparian buffers can help create potential habitat for eels, the density data itself shows places where eels may be present in spite of less than ideal environmental conditions. Another option could be to use CPUE data in future studies, although it has its own drawbacks since it can differ by gear type, which was not listed for ∼75% of samples.

### Recommendations

Based on these results, our recommendations for restoring the American Eel population are to 1) increase access to habitat by re-opening stretches of rivers by removing dams and barriers and 2) increase the quality of the habitat by restoring riparian buffers, which can limit eutrophication from run-off and provide cover for juvenile eels that prefer slower depositional areas with deciduous leaf litter for cover [39].

There are a variety of methods for prioritizing dam removal or fish passage facilities. The Freshwater Network, part of The Nature Conservancy, has built a map of dams organized by aquatic barrier prioritization for the Chesapeake Bay region [51]. A simpler method might be to start with the furthest dam downstream. For example on the Susquehanna River, improving passage at upstream dams without changing the initial dam did not significantly improve upstream eel densities [52]. For migrating salmon, the Washington State Fish & Wildlife Department uses criteria such as habitat suitability, production potential for adults, potential habitat gain, species mobility, species stock condition (whether or not the stock is “of concern” or “depressed”) and potential costs [53, 54]. But since eels have the opposite migration patterns as salmon, this framework may not be completely applicable for conservation measures.

Whether a dam is removed, modified or kept depends on biological, social and economic factors. Dam removal results in hydrologic changes both up and downstream, such as sediment movement and deposition, which alter fine-scale habitat suitability [53]. As such, it may be useful to conduct pre-removal risk assessments, especially in areas where upstream sediments contain pollutants [19]. Dam removal can also be expensive and difficult [55]. Eel ladders are relatively cheap and can provide passage quickly [15] although they may not be ideal for most migratory fish [56]. An eel ladder retrofitted to a dam on the Shenandoah River is accessible to eels between 19 to 74 cm in length. It appears smaller eels are not able to ascend, but most in that size range have probably not yet metamorphosed to yellow eels and therefore are not migrating upstream [57].

An additional benefit to improving access to upstream habitats for eels comes in the distribution of Eastern Elliptio mussels (*Elliptio complanata*). Larvae of freshwater mussels (Bivalvia: Unionidae) are host-dependent and attach to fish hosts until they become free-living juveniles [58]. The mussel, uses American Eel as its primary fish host, but both species are in decline [59]. In the Chesapeake Bay watershed, *E. complanata* recruitment is limited and this appears to be caused by host species distribution, since the mussels are much more abundant downstream of dams on the mainstem of the Susquehanna River than upstream [58]. Restoring American Eel to a stream improves *E. complanata* recruitment but not consistently, since water quality (especially nitrogen and sedimentation) and habitat also play a role [58], which emphasizes the need to improve riparian buffers as well.

Eels can be restocked to areas but this is not a panacea since stocked individuals have different growth rates and sex ratios compared to naturally recruiting eels in the same water body [60] and can carry parasites from one watershed to another [61]. For these reasons we recommend simply re-opening the streams and allowing the eels to recolonize naturally, which has been successful in other watersheds. For example, when the Ft. Edward Dam was removed from the Hudson River, eels were observed in upstream habitats that had been inaccessible for 150 years [62]. Eel abundance also increased significantly after the Embrey Dam in Fredericksburg Virginia was removed in 2004. This dam appeared to have been preventing the migration of smaller individuals and its removal increased eel abundance up to 150 km upstream in less than a year [57].

Additional sampling efforts could provide an opportunity for examining eel morphology across watersheds. Despite the panmixia of the American Eel population (all members can breed with all others), phenotypic differences are evident among different areas in the range, or among different habitats types within a specific region [9].

For future projects, the study area can be expanded to include the entire range of the Chesapeake Bay. The five longest rivers in the Chesapeake Bay watershed are the Susquehanna, Potomac, Rappahannock, York and James rivers are the five largest rivers in the Chesapeake Bay watershed. The map includes the full extent of the latter three rivers but is missing a large portion of the Potomac River and nearly all of the Susquehanna River, which at 715 km is the longest river on the East Coast that drains into the Atlantic Ocean. This, however, will require additional data from Delaware, Pennsylvania, New York and West Virginia. The dataset also includes fish from the Roanoke and Monongahela Rivers (not shown), which are separate drainages.

By adding additional data layers, future projects may be able to address why some eels move upstream and some do not. For example, the map of streams (Fig. 11) could be updated to include flow regimes or eDNA could be used to determine presence/absence in smaller streams that may not have been sampled directly. The methodology in this paper could be used over the entire range of American Eel. For this to be feasible however, there must be standardized data collections from each jurisdiction and an effort to combine multiple datasets into a single database. Future studies will also likely need a different method for categorizing land use, since one category (barren land) barely registered. One option is to use a higher resolution raster to further subdivide land use categories. We initially attempted this work with a different raster set that was available online only, but lost access to this resource (and several other datasets) because of the 2019 U.S. government shutdown. This is what drove us to use DEM data from the ASTER archive (a partnership between the United States and Japan) and land cover data from the Université Catholique de Louvain in Belgium.

Our recommendation to restore habitat and upstream access is supported by analysis by Kahn [12]. This author used data from the National Marine Fisheries to create an index of relative abundance using annual mean total catch of eels per trip, including eels released by anglers (discards), from the period 1981–2014 combined with commercial landings and the index of relative abundance to estimate the trend in commercial fishing mortality in the form of relative fishing mortality. The findings were that the index declined to 1/7^th^ of the original from 1981 to 1995 but increased from 2003 to 2014, although the 2014 index was only about half of what had been observed in 1981. The American Eel fishery has been stable even while abundance has been increasing. For a species with a commercial fishery to be considered endangered with extinction, it would have to become so uncommon as to be commercially extinct, i.e. that a fishery would be economically unviable to the point that landings would not cover the expenses of fishing. Migrating females are less susceptible to the fishery because silver eels tend to stop feeding and are less likely to enter eel pots [12].

### Conclusions

Some subwatersheds, such as the Potomac and Rappahannock Rivers, are of lesser concern, while the areas that merit further study are the York River upstream of Cedar Fork, VA and the James River upstream from Richmond, VA. Both of these watersheds have higher rankings downstream, indicating the high densities of barriers may be affecting upstream eel migration and thus limiting the habitat and carrying capacity of American Eel. Consistent sampling methods and data collection are vital to confirming the results. We recommend removing dams and barriers where appropriate and installing fishways on others, in addition to restoring riparian buffers and improving habitat.

## Acknowledgments

We thank Erik Martin at The Nature Conservancy for advice on studies of fish habitat prioritization. Aaron Bunch at the Virginia Department of Game and Inland Fisheries, Keith Whiteford and Jim Thompson at Maryland Department of Natural Resources and Sheila Eyler at the U.S. Fish and Wildlife Service for help finding biological data. This work was funded by George Mason University as part of the requirements of Walker’s Ph.D. degree.

